# Roost selection by male northern long-eared bats *(Myotis septentrionalis)* in a managed fire-adapted forest

**DOI:** 10.1101/487256

**Authors:** Jesse M. Alston, Ian M. Abernethy, Douglas A. Keinath, Jacob R. Goheen

**Affiliations:** Program in Ecology, Department of Zoology and Physiology, University of Wyoming, Laramie, WY 82071, USA; Wyoming Natural Diversity Database, University of Wyoming, Laramie, WY 82071, USA; Wyoming Ecological Services Field Office, U. S. Fish and Wildlife Service, Cheyenne, WY 82009, USA; Department of Zoology and Physiology, University of Wyoming, Laramie, WY 82071, USA

**Keywords:** Black Hills, Chiroptera, forest management, habitat use, prescribed fire, ponderosa pine *(Pinusponderosa)*, radiotelemetry

## Abstract

Wildlife conservation in multi-use landscapes requires identifying and conserving critical resources that may otherwise be destroyed or degraded by human activity. Summer day-roost sites are critical resources for bats, so conserving roost sites is thus a focus of many bat conservation plans. Studies quantifying day-roost characteristics typically focus on female bats due to their relative importance for reproduction, but large areas of species’ ranges can be occupied predominantly by male bats due to sexual segregation. We used VHF telemetry to identify and characterize summer day-roost selection by male northern long-eared bats *(Myotis septentrionalis)* in a ponderosa pine *(Pinus ponderosa)* forest in South Dakota, USA. We tracked 18 bats to 43 tree roosts and used an information theoretic approach to determine the relative importance of tree- and plot-level characteristics on roost site selection. Bats selected roost trees that were larger in diameter, more decayed, and under denser canopy than other trees available on the landscape. Much like studies of female northern long-eared bats have shown, protecting large-diameter snags within intact forest is important for the conservation of male northern long-eared bats. Unlike female-specific studies, however, many roosts in our study (39.5%) were located in short (≤ 3 m) snags. Protecting short snags may be a low-risk, high-reward strategy for conservation of resources important to male northern long-eared bats. Other tree-roosting bat species in fire-prone forests are likely to benefit from forest management practices that promote these tree characteristics, particularly in high-elevation areas where populations largely consist of males.

## 1. Introduction

Habitat degradation by humans is a leading cause of extinction and population declines of species globally (Dobson et al., 1997; Halpern et al., 2008; Hansen et al., 2013). Less than 15% of Earth’s land surface falls within a protected area, and less than half of that area is free from human development, agriculture, livestock grazing, light pollution, and transportation infrastructure (Jones et al., 2018). Even in relatively undisturbed areas, land uses other than conservation of nature—such as wildfire prevention, livestock grazing, recreation, and extraction of timber and other forest products—are the norm rather than the exception. Conservation measures targeting these multi-use landscapes are thus vital for conserving species (Kremen and Merenlender, 2018).

In multi-use landscapes, successful conservation often requires the identification of critical resources for species of conservation concern so that the supply of those critical resources can be maintained or increased. Day-roosts appear to be critical resources for many bats, providing shelter from predators and environmental stressors (Fenton et al., 1994; Solick and Barclay, 2006), communal sites for social interactions (Willis and Brigham, 2004), and secure places to raise young (Kunz, 1982). Bats spend most of their time in day-roosts, alone or in groups of up to millions of individuals, depending on sex, species, and reproductive status. Patterns of bat abundance and distribution are correlated with roost availability (Humphrey, 1975), and declines in reproductive success have been documented when pregnant or lactating bats are experimentally excluded from preferred roosts (Brigham and Fenton, 1986). Because day-roosts are so important for bats, measures to conserve roosts feature prominently in bat conservation plans. Resource managers seeking to conserve bats while managing landscapes for multiple uses benefit from knowledge that promotes bat roost conservation.

We evaluated day-roost selection by male northern long-eared bats *(Myotis septentrionalis)* in a ponderosa pine forest in the Black Hills of South Dakota, USA. Our study population inhabits an intensively logged landscape at the western edge of this species’ range. Northern long-eared bats inhabit much of the eastern United States and southern Canada (Caceres and Barclay, 2000), but are increasingly threatened by white nose syndrome and have been protected under the Endangered Species Act since 2015. Throughout their range, northern long-eared bats roost almost exclusively in tree cavities and under sloughing bark within intact forest (Lacki et al., 2009), and forage within forests or at forest edges (Henderson and Broders, 2008; Owen et al., 2003; Patriquin and Barclay, 2003).

At our study site and other high elevation areas in the Black Hills, male bats are much more common than females (Choate and Anderson, 1997; Cryan et al., 2000). Sexual segregation driven by elevation or temperature is widespread among bats, and is believed to be driven by differences in energy requirements that allow males to inhabit areas that are colder or have less prey (Barclay, 1991; Ford et al., 2002; Senior et al., 2005). Male northern long-eared bats are therefore likely to occupy substantially different habitat than females, but range-wide conservation for the species is informed predominantly by studies focusing on female bats (J. Alston, unpublished data). Forest managers in male-dominated areas may therefore rely on incomplete information to conserve the majority of bats within their jurisdictions. Our study provides managers in such areas information to appropriately guide management in male-dominated areas and supplement the existing wealth of information on female habitat use.

To evaluate factors driving roost selection, we tracked adult male northern long-eared bats to day-roosts and quantified characteristics of both used and available roost trees using variables easily measured by forest and wildlife managers. We evaluated these data using an information-theoretic approach to select the best models from a suite of candidate models. We hypothesized that in this intensively logged ecosystem, bats primarily select roost trees with characteristics that promote cavity formation (e.g., tree size and amount of decay), the number of nearby roosts (e.g., plot-level tree and snag density), and thermal characteristics suitable for behavioral thermoregulation (e.g., canopy cover and orientation in relation to sunlight).

## 2. Methods

### 2.1 Study Area

We conducted our study during the summers of 2017 and 2018 on Jewel Cave National Monument (43° 45’ N, 103° 45’ W) and surrounding areas of Black Hills National Forest, 16 km west of Custer, South Dakota, USA. In this area, mean monthly summer high temperatures range between 22 – 27° C and mean monthly summer precipitation ranges between 60 – 80 mm (Western Regional Climate Center, 2018). Open ponderosa pine *(Pinus ponderosa)* forests dominate our study site, with Rocky Mountain juniper *(Juniperus scopulorum)* and quaking aspen *(Populus tremuloides)* occurring locally. In our local study area, forests form a heterogenous mosaic with northern mixed-grass prairie where a large stand-replacing fire occurred in 2000. A large cave system and several smaller caves lie underground at our study site, and there is substantial topographic relief on the landscape in the form of intersecting canyon systems and rock outcrops.

Forests in this landscape are intensively managed. Black Hills National Forest typically uses even-aged management techniques other than clear-cutting (e.g., two-step shelterwood harvest). Stand harvest rotations are 120 years on average, but selective cutting occurs at 10-to 20-year intervals to harvest mature trees and thin the understory. Aside from large severe wildfires, the forest self-regenerates and does not require planting. Forest management on private lands generally also follow this formula but thinning intervals vary (B. Phillips, personal communication). Forests on Jewel Cave National Monument are managed for resource preservation, primarily using prescribed fire.

### 2.2 Capture and VHF Telemetry

We used mist nets to capture bats over permanent and semi-permanent water sources (e.g., springs, stock tanks, and stock ponds). In summer (Jun–Aug) 2017 and 2018, we netted 20 and 49 nights at 9 and 15 water sources, respectively. We opened mist nets at civil sunset and closed them after five hours and during inclement weather. We affixed VHF transmitters (LB-2X model .28 g – Holohil Systems Ltd., Carp, ON, Canada; .25 g model – Blackburn, Nacogdoches, TX, USA) between the scapulae of adult male northern long-eared bats with latex surgical adhesive (Osto-Bond, Montreal Ostomy, Montreal, QC, Canada). In our study area and others in the region (Cryan et al. 2000), sex ratios are overwhelmingly male. Because patterns of roost selection can differ between male and female bats (Boland et al., 2009; Elmore et al., 2004; Hein et al., 2008; Perry and Thill, 2007), we targeted males specifically. Additionally, the roosting habits of male bats are less studied than those of females—only 2 of the 14 peer-reviewed studies on roost selection of northern long-eared bats provide data on males, and 11 out of 111 peer-reviewed studies on roost selection of cavity-roosting bats in general provide data on males (J. Alston, unpublished data). All transmitters weighed <5% of the mass of the bat (Aldridge and Brigham, 1988). We tracked bats to roosts each day transmitters were active. All protocols were approved by the University of Wyoming and National Park Service Animal Care and Use Committees and met guidelines approved by the American Society of Mammalogists (Sikes, 2016).

### 2.3 Roost Characterization

To characterize roosts, we collected data for each roost and randomly sampled available roost trees in our study area. We identified available roost trees by generating a sample of 200 random points within 2.53 km (the farthest distance we located a bat roosting from its capture site during our study) of sites where we captured northern long-eared bats and selecting the nearest available roost tree at a random bearing from each point. We defined available roost trees as live trees >20 cm in diameter or any dead tree with a visible defect (e.g. sloughing bark or cavities) sufficiently large for a bat to roost within. For each tree and plot, we measured characteristics that may influence roost suitability (Table 1). We measured vegetation characteristics at two spatial scales: 1) individual trees, and 2) a 706.86 m^2^ (15 m radius) plot around the tree. We also measured topographic variables at the plot scale.

**Table 1.** Variables measured at used and available summer day-roosts of male northern longeared bats *(Myotis septentrionalis)* in the Black Hills of South Dakota, 2017-2018.

| Variable | Definition |
| --- | --- |
| DBH | Tree diameter at breast height (cm) |
| Height | Tree height (m) |
| Snag | Tree status (live/dead) |
| Decay Class | Stage of tree decay on ordinal scale |
| Bark Remaining | Bark remaining on tree stem (%) |
| Canopy Cover | Average of 4 canopy cover measurements (N/E/S/W) taken 5 m from tree (%) |
| Slope | Slope of 706.9 m<sup>2</sup> (15 m radius) plot centered at tree (%) |
| Tree Density | Number of trees in 706.9 m<sup>2</sup> plot centered at tree |
| Snag Density | Number of snags in 706.9 m<sup>2</sup> plot centered at tree |
| Eastness | Difference between aspect of 706.9 m<sup>2</sup> plot centered at tree and 90 degrees (°) |
| Southness | Difference between aspect of 706.9 m<sup>2</sup> plot centered at tree and 180 degrees (°) |
| Slope*Eastness | Interaction term between slope and eastness |
| Slope*Southness | Interaction term between slope and southness |

### 2.4 Statistical Analysis

To quantify differences between roost trees used by northern long-eared bats and the 200 randomly sampled available roost trees, we used the R statistical software environment (R Core Team, 2018) to build binomial-family generalized linear models in a use-availability sampling design (Manly et al., 2007). We employed an information theoretic approach using Akaike’s Information Criterion adjusted for small sample sizes (AIC_c_) to compare competing models (Burnham and Anderson, 2002) using the ‘MuMIn’ package (Barton, 2018). We calculated AIC_c_ values and model weights (w_i_) for all possible combinations of a maximum of 8 predictors (one variable for each 5 observations) in our set of candidate models to prevent bias and unreliable confidence interval coverage (Vittinghoff and McCulloch, 2007). Predictors with variance inflation factors > 10 were removed from consideration in our global model to reduce problems associated with multicollinearity (Kutner, 2005). We averaged model coefficients for all models with cumulative *w_i_* > .95 (Burnham and Anderson, 2002) using the full-averaging method. Finally, we validated our averaged model using area under the receiver operating characteristic curve (AUC; Manel et al., 2001; Swets, 1988).

## 3. Results

We located 44 roosts used on 59 days by 18 bats during our study. Aside from one roost in a rock crevice, bats roosted exclusively in ponderosa pines, either in cavities or under loose bark. Thirty-six out of 43 tree roosts (83.7%) occurred in dead trees (hereafter termed “snags”). We found 2.4 ± 0.3 (range: 1-5) roost trees per bat. Bats typically roosted in the same patch of contiguous forest for the active life of the transmitter. Bats roosted 790 ± 90 m (range: 55 – 2,530 m) from the sites at which they were captured.

Our global model distinguishing used roost trees from available roost trees incorporated DBH, tree height, decay class (Maser et al., 1979), slope, aspect (split into two components— eastness and southness), percent bark remaining, plot tree density, plot snag density, plot canopy cover, and interaction terms between slope and eastness and slope and southness. The global model provided an adequate fit to the data (le Cessie-van Houwelingen-Copas-Hosmer global goodness of fit test; z = 0.805, p = 0.421). Our averaged model (incorporating 104 models in our confidence set; Table A.1) indicated that DBH, decay class, and canopy cover were important variables (Table 2). Significant (p < .05) averaged model coefficients, confidence intervals, and scaled and unscaled odds ratios are reported in Table 3. Mean differences between used and available roost trees among our variables of interest are reported in Table 4. Predictive performance of the averaged model was very high (AUC = 0.924).

**Table 2.**
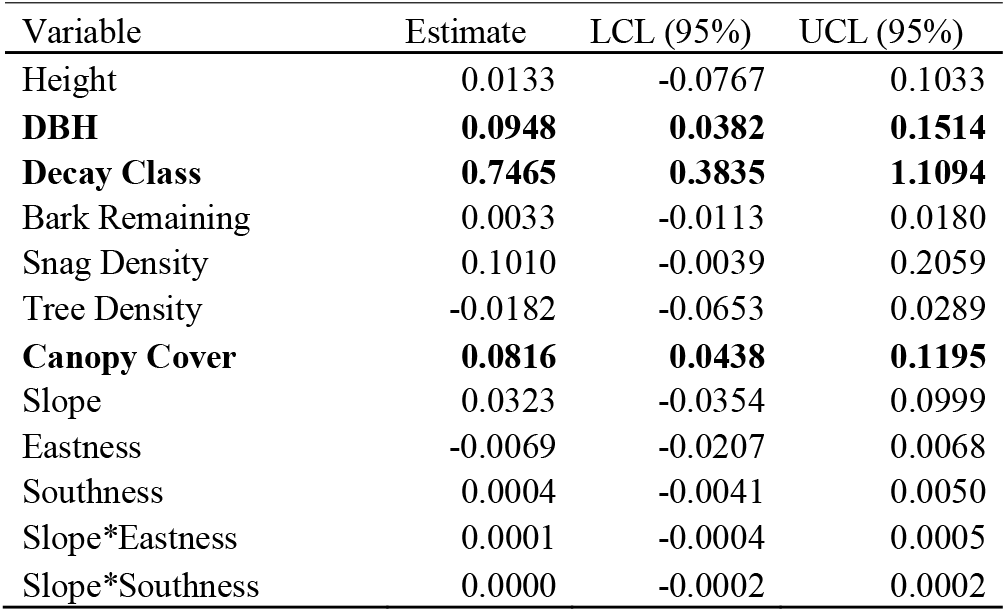
Coefficient estimates in the averaged model and 95% confidence intervals. Bold variables denote significance at *a* = .05.

| Variable | Estimate | LCL (95%) | UCL (95%) |
| --- | --- | --- | --- |
| Height | 0.0133 | -0.0767 | 0.1033 |
| **DBH** | **0.0948** | **0.0382** | **0.1514** |
| **Decay Class** | **0.7465** | **0.3835** | **1.1094** |
| Bark Remaining | 0.0033 | -0.0113 | 0.0180 |
| Snag Density | 0.1010 | -0.0039 | 0.2059 |
| Tree Density | -0.0182 | -0.0653 | 0.0289 |
| **Canopy Cover** | **0.0816** | **0.0438** | **0.1195** |
| Slope | 0.0323 | -0.0354 | 0.0999 |
| Eastness | -0.0069 | -0.0207 | 0.0068 |
| Southness | 0.0004 | -0.0041 | 0.0050 |
| Slope*Eastness | 0.0001 | -0.0004 | 0.0005 |
| Slope*Southness | 0.0000 | -0.0002 | 0.0002 |

**Table 3.** Averaged model coefficients, confidence intervals, and scaled and unscaled odds ratios for significant variables.

| Variable | Coefficient | Unscaled OR | Scaled OR | Units | Scaled OR LCL (95%) | Scaled OR UCL (95%) |
| --- | --- | --- | --- | --- | --- | --- |
| DBH | 0.0948 | 1.0995 | 1.6065 | 5 cm | 1.2105 | 2.1321 |
| Decay Class | 0.7447 | 2.1095 | 2.1095 | 1 unit | 1.4674 | 3.0327 |
| Canopy Cover | 0.0814 | 1.0850 | 2.2619 | 10% | 1.5491 | 3.3025 |

**Table 4.** Means and standard errors for variables of interest among used and available trees. Bold font denotes statistically significant variables in the final averaged model.

| Variable | <u>Roost</u> |  | <u>Available</u> |  |
| --- | --- | --- | --- | --- |
|  | Mean | SE | Mean | SE |
| Height (m) | 8.53 | 1.11 | 9.01 | 0.43 |
| **DBH (cm)** | **35.69** | **1.57** | **30.33** | **0.69** |
| **Decay Class** | **4.95** | **0.33** | **3.72** | **0.18** |
| Bark Remaining (%) | 74.19 | 4.22 | 69.73 | 2.49 |
| Snag Density | 4.74 | 1.03 | 2.12 | 0.23 |
| Tree Density | 19.84 | 2.15 | 10.76 | 1.12 |
| **Canopy Cover (%)** | **36.83** | **3.02** | **14.96** | **1.39** |
| Slope (%) | 16.87 | 1.62 | 11.66 | 0.64 |
| Eastness (°) | 76.36 | 8.21 | 93.35 | 3.81 |
| Southness (°) | 109.48 | 11.14 | 96.58 | 5.48 |

Three variables (DBH, decay class, and canopy cover) were positively related to roost selection (Fig. 1; Table 2). For each 5 cm increase in DBH, odds of selection increased by 61% (CI: 21-113%). For each 1 unit increase in decay class, odds of selection increased by 111% (CI: 47-203%). For each additional 10% increase in canopy cover, the odds of selection increased by 126% (CI: 55-230%).

**Fig. 1.**
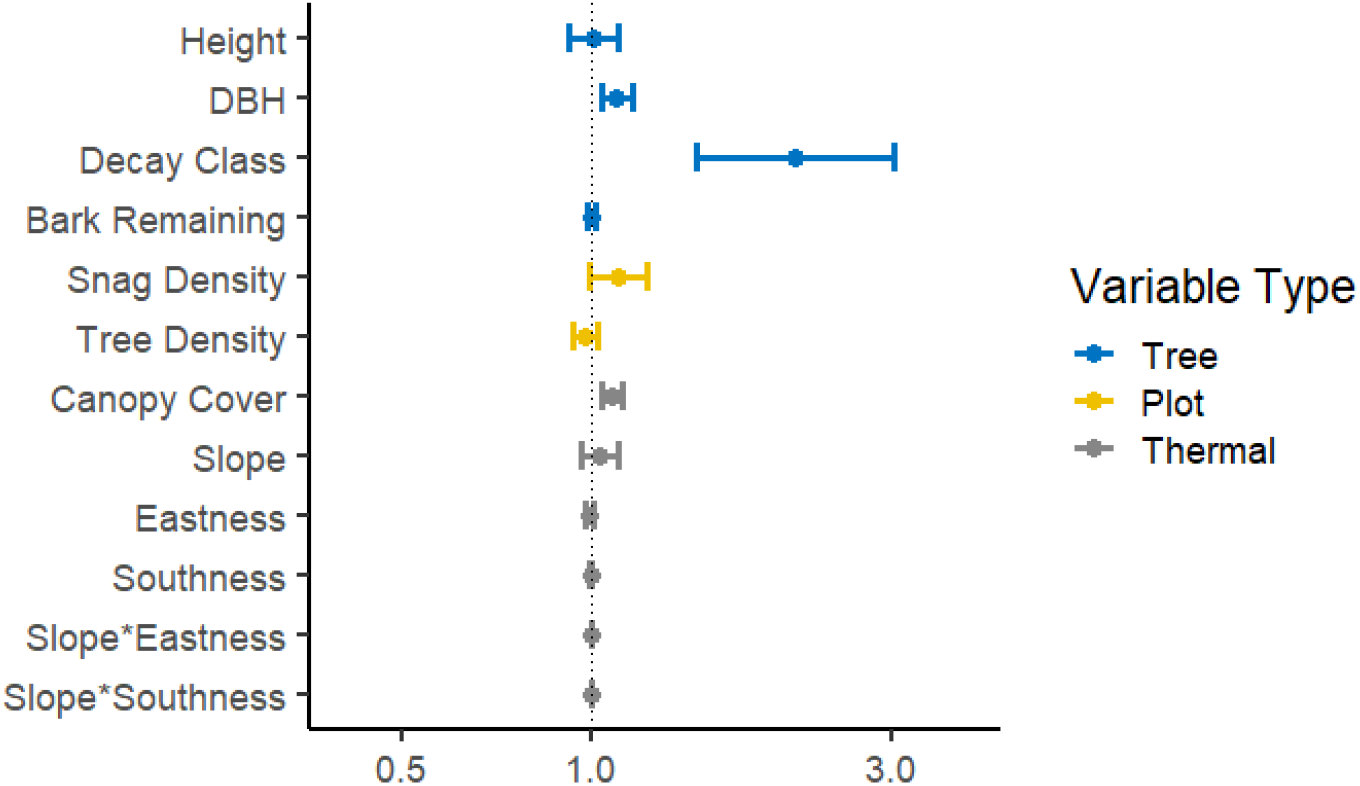
Unscaled odds ratios associated with each variable in the averaged roost selection model. Error bars represent 95% confidence intervals.

## 4. Discussion

Northern long-eared bats primarily selected roosts in trees with characteristics that promote cavity formation. At the level of individual trees, northern long-eared bats selected for large diameter trees with substantial decay. This corroborates previous work on northern long-eared bats (Jung et al., 2004; Rojas et al., 2017) and is intuitive because large trees with more decay have more roost structures (i.e., cavities and loose bark) for bats to use (Reynolds et al., 1985). This is particularly true of ponderosa pines, which can produce large amounts of resin to defend against physical injury (Kane and Kolb, 2010; Lewinsohn et al., 1991) and therefore tend to develop cavities only when they are scarred or dead. In intensively logged landscapes like the Black Hills, cavities are found overwhelmingly in snags because most trees are harvested before they reach ages at which cavities typically form.

Conservation actions targeting northern long-eared bats should include preservation of large snags whenever possible. Our study demonstrated that northern long-eared bats select large-diameter snags (>37 cm), and large diameter snags also tend to remain standing longer than thinner snags (Bull, 1983; Chambers and Mast, 2014). These snags need not be tall—short (≤ 3 m) snags are important resources for male northern long-eared bats as well. Seventeen of 43 (39.5%) roosts that we located occurred in broken-off snags ≤ 3 m in height. These are important resources and are likely more vulnerable to loss during forest management activities (particularly prescribed fire) than other potential roost trees. Snags are often intentionally removed during forest management activities because of hazards posed to forest management personnel (e.g., loggers and firefighters) and the general public. However, these short snags pose less danger to forest management personnel and the public than taller snags, and their preservation is therefore a realistic and actionable step toward bat conservation.

Of the variables we considered that may influence thermal characteristics of roosts, only canopy cover influenced roost selection significantly. Trees were more likely to be used as roosts as surrounding canopy cover increased, and use was greater than availability at all canopy cover levels >19%. Although many snags were available at our study site in open areas burned by a severe wildfire in 2000, northern long-eared bats rarely use those snags, instead selecting snags in the interior of forest stands with live canopy. Forty out of 43 (93.0%) roosts were within or immediately bordering intact forest stands with live canopy, and all roosts were within 50 m of intact forest stands. Bats may prefer these areas because canopy cover creates cooler environments, but they may also simply prefer to be immediately near forested areas where they forage (Henderson and Broders, 2008; Owen et al., 2003; Patriquin and Barclay, 2003). Either way, stand-replacing fire likely poses risks to local populations of northern long-eared bats at the western edge of its range, where severe wildfire is increasingly prevalent due to climate change (Westerling et al., 2006). Clearcutting also poses risks to local populations of northern long-eared bats in these areas, even if snags are retained. Selective logging that leaves some level of canopy cover remaining would ensure that snag retention is effective for bat roost conservation.

Dynamics of regional disturbance may be important when evaluating local-scale factors that influence roost selection (O’Keefe and Loeb, 2017). The ponderosa-dominated landscape where we conducted our research is substantially different than other landscapes (i.e., deciduous and mixed forests in eastern North America) where roost selection by northern long-eared bats has been studied. Although many of the factors driving roost selection appear to be similar among areas, the processes that create roosts may be fundamentally different in different areas. Snags in ponderosa pine forests are often generated in large pulses by severe wildfire and mountain pine beetles *(Dendroctonus ponderosae)*, but the long-term ramifications of these resource pulses for bats are not well understood. Severe wildfire appears to create snags that are largely unused by bats. Mountain pine beetle outbreaks may do the same if beetle-induced mortality reduces or eliminates canopy cover over large areas, or if outbreaks lead to more severe fires. Northern long-eared bats may instead depend on snag-generating processes that operate at more local scales and over longer intervals to create suitable roosts.

Roost selection by bats varies by sex, age class, and reproductive condition (Elmore et al., 2004; Hein et al., 2008). Studies on roost selection generally focus on females because they tend to drive reproduction, which is required to sustain populations. However, targeting roost conservation toward females exclusively may neglect resources that are important for males. Because sex ratios can be heavily biased in some areas (Cryan et al., 2000), ignoring the needs of males could leave resources that are important for most individuals inhabiting these areas unprotected. On the other hand, designing roost conservation measures on studies of males alone will leave resources that are important for females unprotected. For example, short (≤ 3 m) snags are important resources for males, but they may not be for females, which aggregate in maternity colonies that require larger cavities than largely solitary males (Perry and Thill, 2007). Resource managers seeking to conserve bats should take these sex differences into account when developing conservation plans and designing studies to inform those plans. In high elevation areas, males may be more important than females for sustaining local populations because there are few females in those areas.

## 5. Conclusions

Forest managers require actionable knowledge to guide conservation, and our results indicate that conserving large-diameter snags within intact forest stands is one such action that can be taken to conserve bats in wildfire-prone coniferous forests. Short (≤ 3 m) snags in particular represent a low-risk, high-reward resource to target for preservation in male-biased, high elevation populations. For federally threatened northern long-eared bats, conserving these snags at the western edge of their range may prevent range contraction and local extinction. Similar patterns are likely to hold true for other cavity-roosting bat species in wildfire-prone coniferous forests, like those found throughout western North America. Although bats face danger from many threats unrelated to roosts (e.g., white nose syndrome, wind energy development, etc.), roost conservation remains an important tool for bat conservation in the face of such threats.

## Acknowledgements

Many thanks to L. Boodoo, C. McFarland, E. Greene, B. Tabor, and B. Phillips for help with fieldwork; J. Rick for helpful comments on pre-submission versions of this manuscript; R. Anderson-Sprecher for helpful comments concerning statistical analyses; and P. Ortegon, D. Licht, M. Wiles, D. Austin, B. Phillips, and E. Thomas for their logistical support of this project. Research funding was provided by the National Park Service, the Department of Zoology and Physiology at the University of Wyoming, the Berry Ecology Center, the American Society of Mammalogists, Prairie Biotic Research, Inc., and the Wyoming Chapter of The Wildlife Society. The findings and conclusions in this article are those of the authors and do not necessarily represent the views of the U.S. Fish and Wildlife Service or the National Park Service.

## Appendix A: Supplementary Data

**Table A.1**. Candidate models, ΔAIC values, and model weights (*w_i_*) used to determine model-averaged coefficients.

## Data Availability

*Data and R code used in analysis are available in the supplementary material and will be archived on the lead author’s personal website if this manuscript is accepted for publication.

